# A metagenomic framework for rapid *Listeria monocytogenes* surveillance in food production environments

**DOI:** 10.64898/2026.04.23.720354

**Authors:** Francis Muchaamba, Tim Reska, Michael Biggel, Kristopher M. Locken, Lukas Weilguny, Sabrina Corti, Lucien Kelbert, Roger Stephan, Lara Urban

## Abstract

*Listeria monocytogenes* remains a major foodborne pathogen with high mortality and costly persistence in food-processing environments. Established diagnostics rely on selective enrichment and single-colony isolation, which could introduce strong biases by favouring fast-growing strains or those more tolerant to enrichment broth inhibitors, while suppressing slow-growing, viable-but-nonculturable, and other co-occurring strains. This can obscure true pathogen diversity and may contribute to discrepancies between strains detected in food production environments and those associated with disease. To quantify the bias introduced by established culture-based diagnostics and to assess the potential advantage of metagenomics-based pathogen detection directly from the original sample matrix, we developed and evaluated a rapid nanopore sequencing–based metagenomic framework. We designed an artificial metagenomic community of several *Listeria* strains, comprising *L. monocytogenes* lineages I–III (including hypervirulent, persistent, and low-virulence strains), other *Listeria* spp., and a realistic background microbiome representative of food-processing environments. We then used this mock community to spike standard surveillance sponges and compared three workflows: (*i*) direct nanopore metagenomic sequencing of the original sample matrix, (*ii*) quasi-metagenomic sequencing after 4 h, 12 h, 24 h, or 48 h of selective enrichment, and (*iii*) ISO-based culture followed by whole-genome sequencing of a single presumptive *L. monocytogenes* isolate. We found that the culture-based approach recovered only a limited subset of strains, consistently underrepresenting diversity and failing to detect multi-strain contamination. These findings were reflected by the quasi-metagenomic results, where we found relative *L. monocytogenes* enrichment to be strain-dependent, indicating selective enrichment bias favouring specific strains. Metagenomics captured the full spectrum of spiked *Listeria* strains, enabling comprehensive strain-level resolution at all inoculation levels. We only observed relative enrichment of the *L. monocytogenes* strains by quasi-metagenomics compared with metagenomics after 48 h of selective enrichment. While driven primarily by the additional enrichment of *L. innocua*, these results suggest that quasi-metagenomics improves *L. monocytogenes* recovery only at the cost of a substantial reduction in speed. We finally showed that the sensitivity and accuracy of metagenomics could be improved by utilising different environmental sampling materials. We did not find any significant performance improvements from nanopore sequencing-based enrichment of *L. monocytogenes* through adaptive sampling approaches. We conclude that integrating long-read metagenomics into routine surveillance shows great promise to improve detection and source attribution in food safety systems.

## Introduction

*Listeria monocytogenes* remains one of the most consequential foodborne pathogens, responsible for severe invasive disease and disproportionately high mortality (European Food Safety Authority (EFSA) and European Centre for Disease Prevention and Control (ECDC), 2025). Its ability to persist in food-processing environments, survive in multispecies biofilms, and withstand sanitation regimes poses ongoing challenges for the food industry and public health systems (Finn et al., 2023; Osek et al., 2022). Despite extensive prevention and control measures and regulatory oversight, contamination events continue to occur, often involving complex mixtures of strains that vary in virulence, stress tolerance, and environmental persistence (Muchaamba et al., 2022). The *L. monocytogenes* genotypes most frequently isolated from foods and food-processing environments are stress-tolerant, weakly virulent clonal complexes such as CC9 and CC121, which are not the genotypes most commonly associated with clinical listeriosis (Nüesch-Inderbinen et al., 2026; Maury et al., 2019; Maury et al., 2016). Instead, hypervirulent lineage I strains, particularly serotype 4b (CC1, CC4, and CC6 strains), are disproportionately represented among human cases (Painset et al., 2019). This might suggest that current surveillance methods do not accurately reflect the *L. monocytogenes* populations present in foods and food-processing environments.

Established diagnostics rely almost entirely on selective enrichment followed by the characterisation of a single presumptive colony (ISO 11290-1). Although this approach offers practicality and standardisation, it imposes strong selective pressures that might distort the underlying population structure of pathogens and potentially obscure the strains of epidemiological importance. Fast-growing strains or those more tolerant to enrichment broth inhibitors (such as, e.g., acriflavine, lithium chloride) are preferentially multiplied, while slow-growing or otherwise disadvantaged (viable but non-culturable (VNBC) or L-form) strains are frequently suppressed to non-detectable levels (Zilelidou et al., 2016; Bruhn et al., 2005). Consequently, culture-based diagnostics often yield an artificial representation of the population present *in situ*. The routine practice of single-colony selection further tightens this bottleneck, effectively precluding the detection of multi-strain contamination, which is, however, increasingly observed in food and food production environments: For example, recent work in *Vibrio parahaemolyticus* has highlighted the extent of this issue, revealing unexpectedly high within-matrix strain diversity, with more than ten genetically distinct sequence types recovered from a single seafood sample.

Culture-independent metagenomic approaches offer the potential to overcome these limitations by enabling rapid, unbiased detection and strain-level characterisation of pathogens directly from any input matrix and therefore capturing strains that might be lost during enrichment (Walsh et al., 2017; Wang et al., 2025), including rare, novel, or VBNC pathogens (Flörl et al., 2025; Ghiotto et al., 2024). In addition to pathogen identity, metagenomics can simultaneously resolve virulence potential, antimicrobial resistance determinants, and the distribution of mobile genetic elements such as plasmids (Ürel et al., 2026). Unlike targeted approaches, it can further detect emergent diagnostic escapes and distinguish them from true reductions in transmission. While the power of strain-resolved metagenomics has already been demonstrated in real-world outbreak settings (Buytaers et al., 2021), the low microbial biomass typically present in environmental samples often restricts the sensitivity and resolution of metagenomic approaches. The following laboratory and genomic approaches have been suggested to improve metagenomic performance: In the laboratory, DNA extraction optimisations can significantly increase yield; alternatively, quasi-metagenomic approaches apply metagenomics after minimal selective enrichment, which can enhance pathogen recovery while avoiding the severe selective distortions inherent to established culture-based diagnostics (Kocurek et al., 2023; Wagner 2021). For analytical optimisations, long-read sequencing as provided by the nanopore sequencing technology has been shown to offer increased strain-level resolution and pathogen recovery (Bellankimath et al., 2026; Ürel et al., 2026; Tan et al., 2025; Urban et al., 2023); in addition, nanopore sequencing allows for selective enrichment or depletion of genomic material based on reference sequences using so-called “adaptive sampling” during sequencing (Martin et al., 2023; Loose et al., 2016), which might enable the selective enrichment of pathogens of interest without additional sample manipulation.

To quantify the bias introduced by established culture-based diagnostics and to assess the potential advantage of metagenomics-based pathogen detection directly from the original sample matrix, we developed and evaluated a rapid nanopore sequencing–based metagenomic framework using realistic microbial mock communities. We implemented laboratory and genomic optimisations to assess whether direct metagenomics could improve sensitivity, speed, and strain-level resolution compared with established culture-based diagnostics, and to assess the distortion in the detection and representation of *L. monocytogenes* introduced by selective enrichment and single-colony characterisation.

## Results

Using a mock microbial community of several *L. monocytogenes* strains and other *Listeria* species at different concentrations, together with a realistic environmental background microbiome, we compared metagenomic, quasi-metagenomic, and standard culture-based approaches for *L. monocytogenes* strain detection (**Figure 1**; Methods).

**Figure 1.**
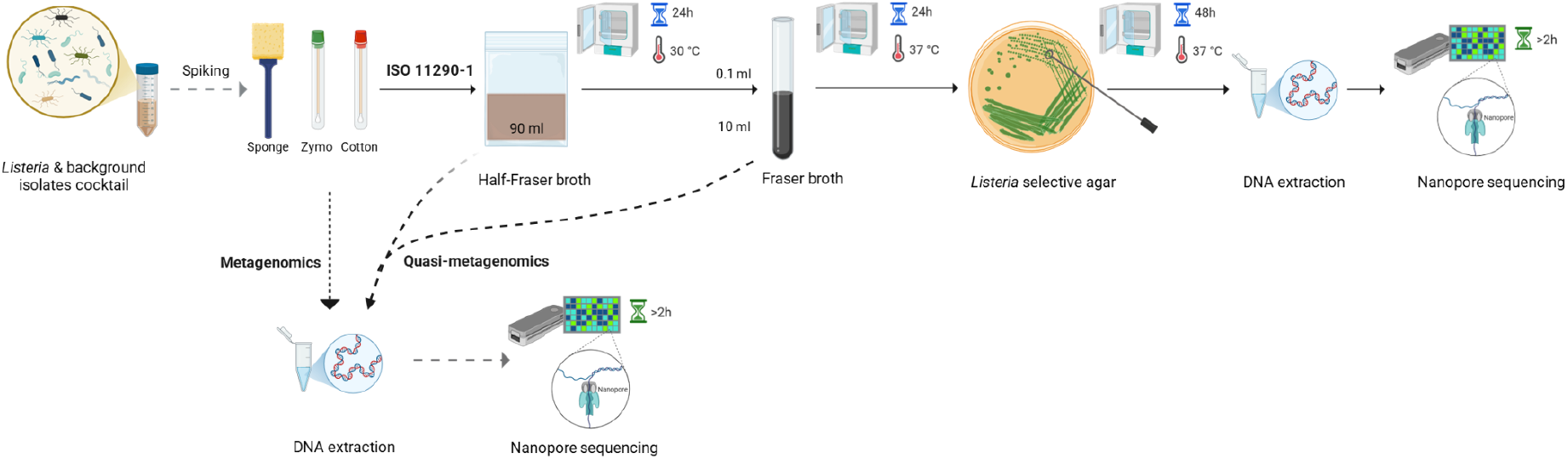
Comparison of metagenomics, quasi-metagenomics, and culture-based diagnostics by whole-genome sequencing for *Listeria monocytogenes* detection. Environmental sampling devices were artificially contaminated with a defined *L. monocytogenes*, other *Listeria* species, and a background microbiome. Samples were processed according to ISO 11290-1, including enrichment in Half-Fraser broth (HFB; 24 h at 30 °C), followed by Fraser broth (FB; 24 h at 37 °C) and plating on selective agar (48 h at 37 °C). DNA extraction and nanopore sequencing were performed at three stages: (i) directly from samples (metagenomics: dotted arrow), (ii) after HFB and FB enrichment (quasi-metagenomics: dashed arrows), and (iii) from presumptive *L. monocytogenes* colonies recovered from selective agar (ISO 11290-1 approach: solid arrows). Created with BioRender.

We first used sponges as the current reference matrix for environmental surveillance to compare metagenomic, quasi-metagenomic, and culture-based approaches (Methods). While DNA concentrations increased with enrichment time (especially after 24 h of enrichment) during quasi-metagenomics, the relative abundance of *L. monocytogenes* increased compared with metagenomics only after 48 h of enrichment (**Figure 2**). Adaptive sampling did not significantly increase the relative abundance of *L. monocytogenes* across all time points (**Figure 2**).

**Figure 2.**
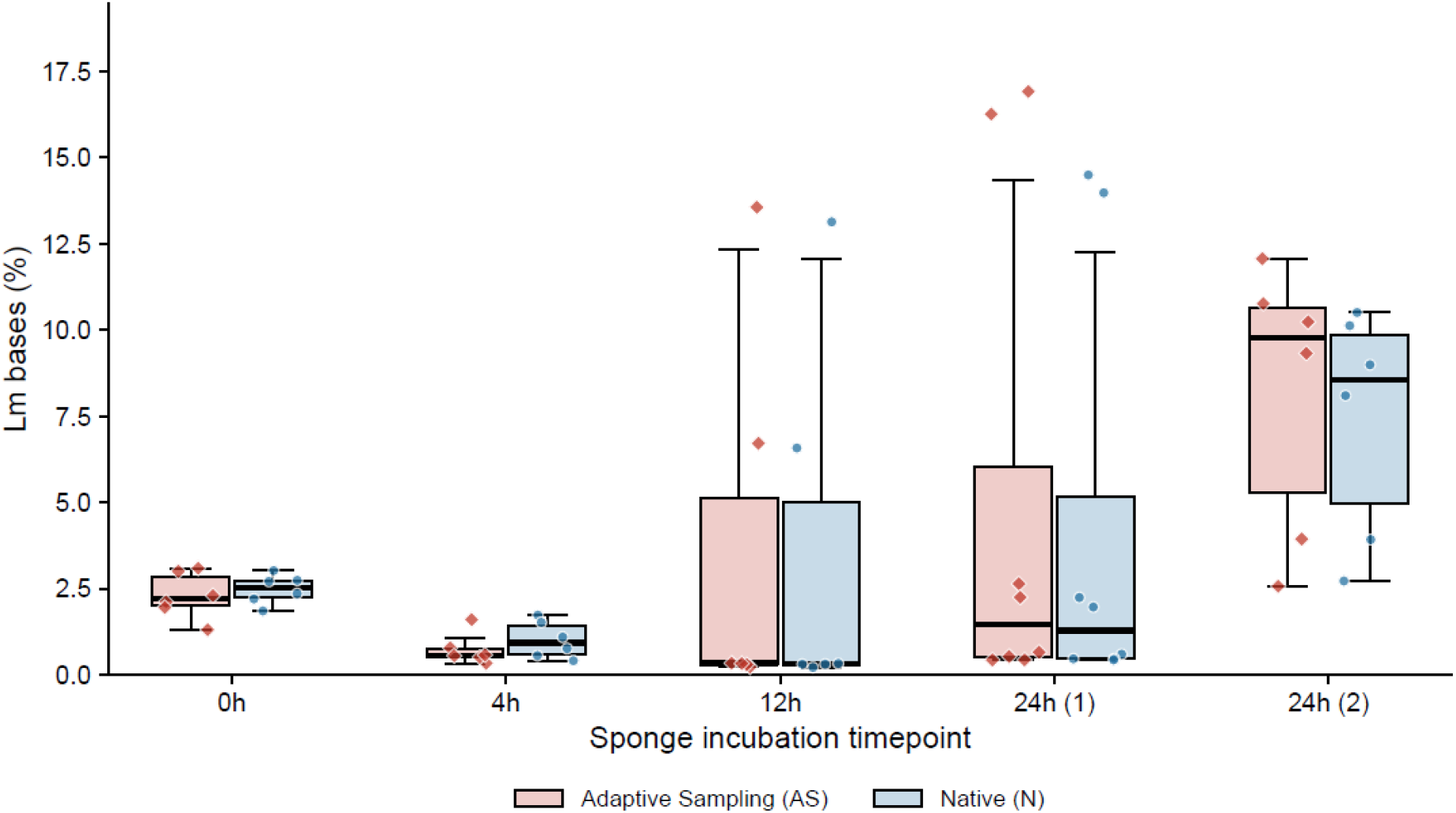
Relative abundance of *L. monocytogenes* (Lm) bases (%) in sponges in the quasi-metagenomic dataset (after selective enrichment: 4 h, 12 h, 24 h in HFB, and an additional 24h in FB (“24h (2)”)) in comparison to the metagenomic dataset (0h), and in comparing the adaptive sampling (AS) and normal (N) nanopore sequencing mode. Points represent biological replicates; boxplots show median and interquartile range.

Based on the standard nanopore sequencing data (no adaptive sampling), metagenomic sequencing of the sponge samples detected all spiked *L. monocytogenes* strains across all inoculum levels (Lm2, Lm4, Lm6; **Figure 2**: x=0h; **Supplementary Table S1**). Among the four *L. monocytogenes* strains analysed, at t = 0h LMNC326 was consistently detected as the most abundant strain (**Figure 3**). Detection sensitivity at the lowest inoculum level (Lm2) was sufficient to recover *L. monocytogenes* reads across all replicates, although relative abundance remained low compared to higher inoculum levels (**Supplementary Table S1**). Quasi-metagenomic sequencing also detected all *L. monocytogenes* strains at all time points, but relative abundance profiles indicated increasing dominance of specific genotypes over time (**Figure 3**; **Supplementary Table S2**): LL195 (ST1) and N13-0119 (ST121) showed a consistent increase in relative abundance over time, whereas EGDe (ST35) and LMNC326 (ST70) did not (**Figure 3**).

**Figure 3.**
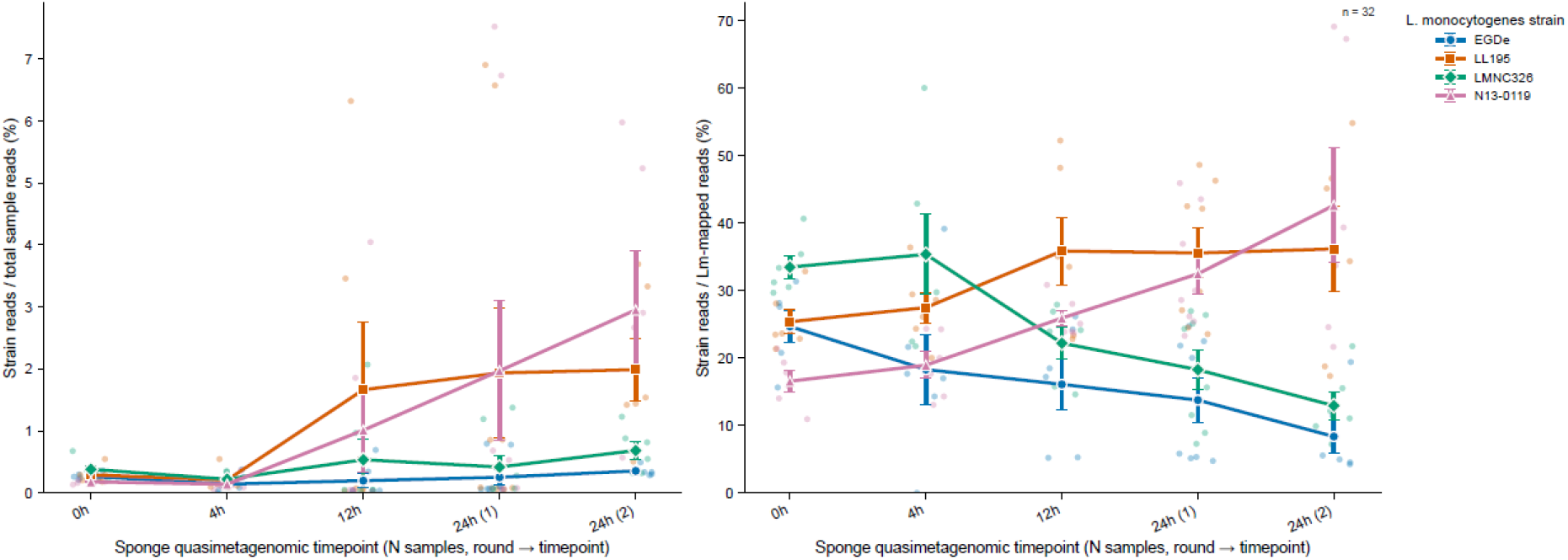
Relative abundance of *L. monocytogenes* (Lm) strains (%) in sponges in the quasi-metagenomic dataset across all inoculum concentrations (after selective enrichment: 4 h, 12 h, 24 h in HFB, and an additional 24h in FB (24h (2))) in comparison to the metagenomic dataset (0h). Semi-transparent points represent biological replicates; solid points represent the mean; and error bars represent the Standard Error of the Mean.

At the *Listeria* species level, similar patterns were observed (**Figure 4**): While the metagenomic datasets detected all four *Listeria* species (*L. monocytogenes, L. innocua, L. welshimeri*, and *L. ivanovii*) at relative abundances consistent with the defined input composition, *L. innocua* and *L. monocytogenes* consistently increased in relative abundance throughout the selective enrichment process, while *L. welshimeri* and *L. ivanovii* were partially or completely outcompeted.

**Figure 4.**
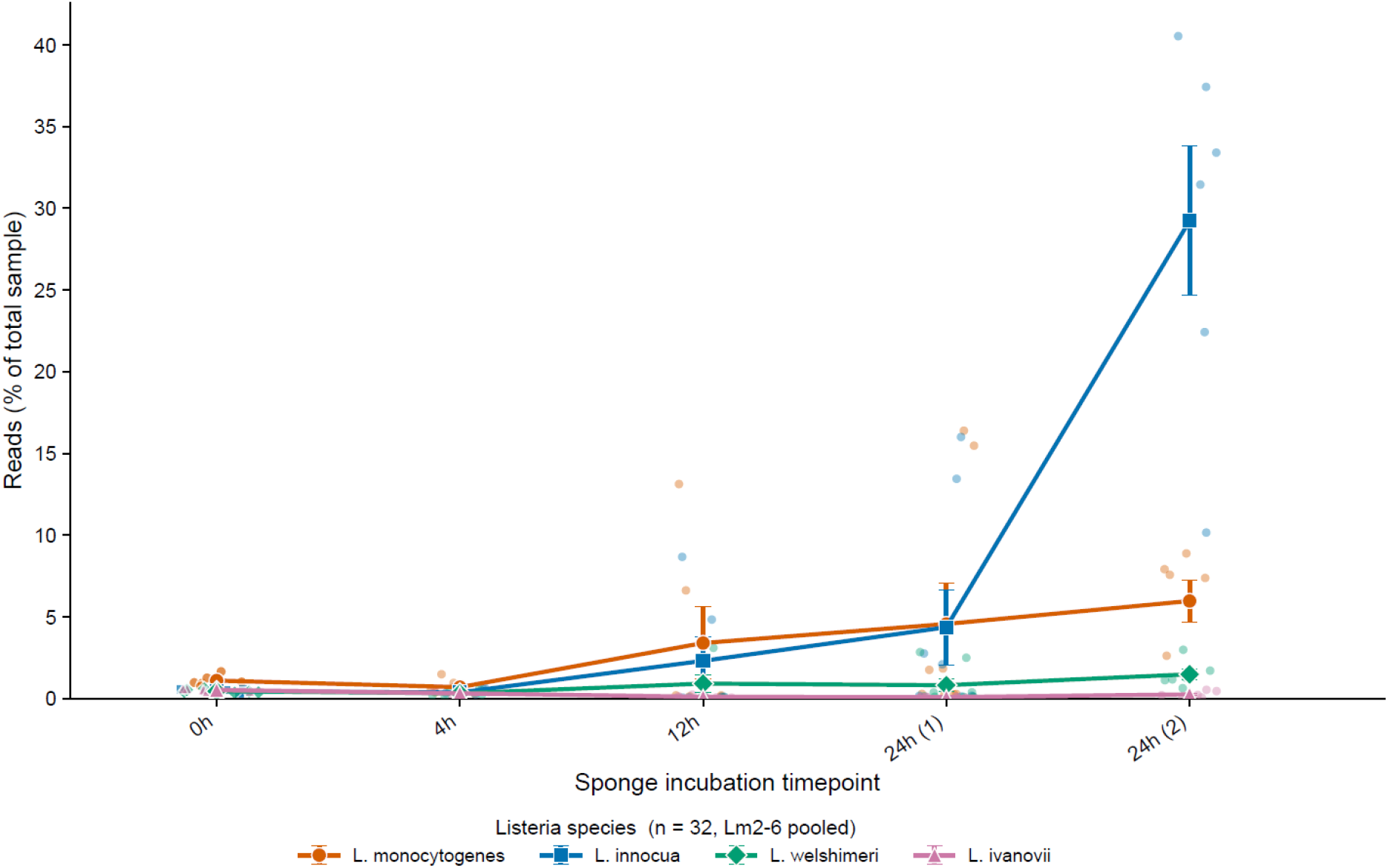
Relative abundance of all *Listeria* species (%) in sponges in the quasi-metagenomic dataset (after selective enrichment: 4 h, 12 h, 24 h in HFB, and an additional 24 h in FB (24h (2))) compared with the metagenomic dataset (0h). Semi-transparent points represent biological replicates; solid points represent the mean; and error bars represent the Standard Error of the Mean.

In contrast to the metagenomic and quasi-metagenomic datasets, which were generated from mixed-community samples, whole-genome sequencing was performed only on single presumptive *L. monocytogenes* isolates recovered by the established culture-based workflow (**Supplementary Table S3-S4**). This isolate-based whole-genome sequencing consistently identified *L. monocytogenes*, but recovered only a subset of the strains present in the inoculum, thereby underrepresenting overall diversity and failing to capture co-occurring strains. Among the four spiked *L. monocytogenes* strains (Methods), LL195 (ST1) and N13-0119 (ST121) were consistently recovered, whereas EGDe (ST35) and LMNC326 (ST70) were not detected (**Supplementary Table S4**). This observation agrees with the consistent relative increase of the ST1 and ST121 strains observed in the quasi-metagenomic data (**Figure 3**).

Despite the robust recovery of spiked pathogens from the sponge metagenomic data, we observed substantial biomass retention within the sampling matrix in comparison with standard cotton and Zymo swabs (**Supplementary Tables S1** and **S2**). The relative abundance of *L. monocytogenes* reads varied across the three sampling matrices: Sponges showed the lowest relative abundance, while cotton swabs yielded the highest relative abundance and most consistent detection across all spiking levels (**Figure 5**). These results indicate that the sampling matrix influences both DNA recovery and downstream detection performance in metagenomic workflows.

**Figure 5.**
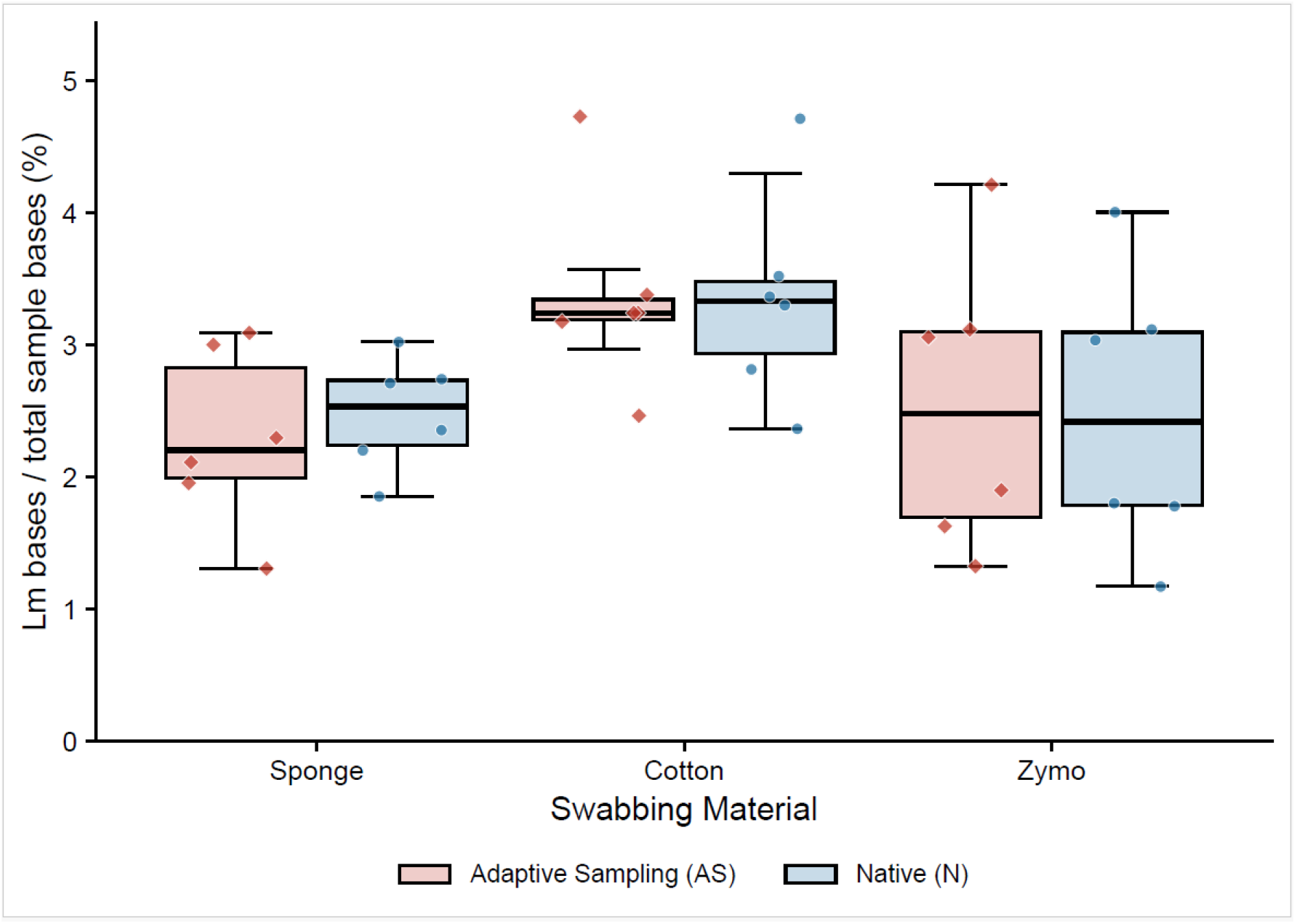
Effect of sampling matrix on relative abundance of *L. monocytogenes* (Lm) bases (%) in sponges, cotton, and Zymo swabs, comparing adaptive sampling (AS) and normal (N) nanopore sequencing modes. Points represent biological replicates; boxplots show median and interquartile range.

## Discussion

In this study, we developed and systematically evaluated a rapid nanopore sequencing–based metagenomic framework using defined, multi-strain mock communities, and directly compared its performance to established selective enrichment and culture-based workflows. This experimental design enabled controlled assessment of detection sensitivity, strain recovery, and enrichment-induced bias across workflows.

We found that selective enrichment and culture-based approaches substantially distort the original strain composition and consistently underrepresent *L. monocytogenes* strain diversity by failing to capture multi-strain contamination. We further demonstrate that enrichment reproducibly favours specific strains across all inoculum levels, resulting in a divergence between the original population structure and the strains recovered after cultivation. This confirms longstanding concerns that reliance on single-colony isolation and selective enrichment introduces substantial bias, preferentially detecting fast-growing strains or strains with increased tolerance to selective agents, e.g., acriflavine or lithium chloride (Zilelidou et al., 2016; Bruhn et al., 2005; Suh and Knabel, 2001; Jacobsen, 1999).

Consequently, culture-based workflows systematically underestimate strain diversity and may misrepresent mixed contamination events. At the species level, enrichment produced similar distortions. While the metagenomic data accurately represented the inoculum composition, selective enrichment resulted in reduced diversity and increasing dominance of *L. monocytogenes* and *L. innocua*, whereas *L. welshimeri* and *L. ivanovii* declined to low abundance or became undetectable. These results demonstrate that enrichment does not merely enhance detection sensitivity but actively reshapes community structure, introducing both strain- and species-level bias. Notably, *L. innocua* consistently outcompeted other *Listeria* species, including *L. monocytogenes*, under the tested conditions. These observations might suggest that *L. monocytogenes* detections can be hampered by the presence of *L. innocua*, which has been reported to be a potential ecological indicator of *L. monocytogenes* (Milillo et al., 2012; Manqele et al., 2023). Similar impact of *L. innocua* and other *Listeria* spp. on *L. monocytogenes* detection has been reported (Dailey et al., 2015; Zitz et al., 2011; Cornu et al., 2002; Scotter et al., 2001). Of note, only a single *Listeria* enrichment workflow (HFB and FB) was evaluated in this study; alternative enrichment media and protocols may impose different selective pressures and could result in distinct strain- and species-level dynamics, which should be assessed in future work.

A closer examination of enrichment dynamics reported by Ottesen et al. (2020) provides additional context for interpreting quasi-metagenomic data (Ottesen et al., 2020). In their study, *L. monocytogenes* remained at low relative abundance (∼1–2% of reads) at 20 h of enrichment, and increased to approximately 15% by 24 h, representing a rapid shift in community composition over a short time interval. While this increase provided sufficient genomic coverage for outbreak-level phylogenetic analysis, it also indicates a strong and selective expansion of *L. monocytogenes* relative to the surrounding microbiota. However, the implications of this shift for within-species diversity were not explored. Our results demonstrate that such rapid increases in relative abundance are accompanied by pronounced strain-level selection, with specific genotypes consistently outcompeting others during enrichment. Notably, we observe that compositional changes begin even earlier (4–12 h), suggesting that the underlying population structure may already have diverged substantially from the original sample at 20-24 h. While quasi-metagenomic sequencing at this time point appears to accurately identify dominant strains (Ottesen et al., 2020), it may underrepresent co-occurring or less competitive lineages, highlighting an inherent limitation of enrichment-based approaches for resolving true within-sample diversity.

Metagenomics, on the other hand, enabled culture-independent detection and strain-level profiling of all *L. monocytogenes* strains and all other spiked *Listeria* species in our study. It is, however, important to note that strain-level identification in this study was based on mapping of sequencing reads against reference genomes of the spiked strains, and therefore represents a reference-guided approach; the performance of fully reference-free or *de novo* strain resolution was not yet assessed. While quasi-metagenomics improved DNA yield, an important advantage for low-biomass samples, the relative abundance of enriched *L. monocytogenes* only surpassed the original metagenomic samples after 48 h of enrichment. This time requirement, together with the reliance of quasi-metagenomics on biosafety spaces, might make it less accessible and applicable for rapid pathogen surveillance. Rapid identification of foodborne pathogens is, however, essential for timely outbreak mitigation, reducing disease burden, and preventing the distribution of contaminated food. Similar to metagenomics, quasi-metagenomics also recovered all *L. monocytogenes* strains. However, we found that selective enrichment before final culturing introduced a substantial taxonomic compositional shift, probably contributing to the biased culture-based detections. Specifically, only two *L. monocytogenes* strains, namely LL195 (ST1) and N13-0119 (ST121), could be recovered from the culture-based diagnostics of all samples, and the same two strains consistently increased in relative abundance throughout the selective enrichment protocol.

Our findings extend recent applications of metagenomics in food safety surveillance. The first large-scale food production metagenomic study (Quijada et al., 2025) applied short-read sequencing to characterise microbial communities and resistomes across food environments (Barcenilla et al., 2024). While informative, short-read approaches remain limited in their ability to resolve strain-level diversity and to improve inference of mobile genetic elements, such as plasmids, in relation to their microbial hosts. In contrast, nanopore-based long-read metagenomics provides improved genomic contiguity, enabling higher-resolution strain reconstruction, more robust host–plasmid association, and access to additional epigenetic signals such as DNA methylation patterns (Ürel et al., 2026).

In addition to sequencing technology, the sampling strategy represents a key determinant of metagenomic performance. We found that sponges exhibited partial retention of bacteria, resulting in reduced recoverable DNA for metagenomic sequencing. More generally, DNA recovery in environmental sampling appears to be strongly influenced by the physical and chemical properties of the sampling matrix, including porosity and biomass retention characteristics. This suggests that sampling materials should not be treated as interchangeable, but rather as a key design parameter that influences metagenomic sensitivity. While pre-moistened sponges remain widely used in food safety workflows, alternative matrix compositions may offer improved DNA release efficiency and more consistent recovery, depending on downstream analytical requirements. Consistent with this, swab-based sampling yielded higher amounts of recoverable DNA and of *L. monocytogenes* genomic information in our setup. While this approach slightly increased our sensitivity, nanopore sequencing-based enrichment through adaptive sampling did not have an impact. This finding might be explained by the relatively short sequencing reads (median read length of 1,093 bases) in our metagenomic dataset, which can make adaptive sampling less efficient, since no substantial advantage is gained from rejecting sequencing reads, while the read rejection leads to faster pore usage and sequencing lag time. Our DNA extraction approach should therefore be further optimised to obtain longer reads and potentially improve adaptive sampling efficiency in future applications.

In summary, our findings demonstrate that culture-independent approaches can complement established diagnostics by improving strain-level resolution and reducing enrichment-associated bias. The combined use of optimised sampling, extraction, and sequencing strategies has the potential to enhance detection accuracy and provide a more comprehensive understanding of *L. monocytogenes* populations in environmental monitoring contexts. The inability of culture-based workflows to capture full strain diversity may obscure mixed contamination events and hinder accurate source attribution. This is particularly relevant for *L. monocytogenes*, where strain-level differences are linked to stress tolerance and virulence potential. These findings have direct implications for food safety surveillance, where accurate assessment of strain diversity and low-abundance contaminants is critical for outbreak detection and source attribution. More broadly, these findings might support a shift toward integrating strain-resolved metagenomic approaches into routine surveillance frameworks and environmental monitoring. By rapidly capturing the complete genomic landscape directly from a sample, metagenomics enables simultaneous characterisation of pathogen identity, diversity, and potential functional traits, including virulence and antimicrobial resistance determinants (Bellankimath et al., 2026; Ürel et al., 2026; Parks et al., 2026; Olsen and Riber, 2025; Liu et al., 2023). This represents a substantial advancement over traditional single-isolate approaches, which inherently limit resolution and may overlook clinically relevant variants. Continued advances in sequencing technologies, enrichment strategies, and bioinformatic pipelines will further enhance the applicability of these approaches in real-world settings.

## Acknowledgements

We thank the staff at the Institute for Food Safety and Hygiene, University of Zurich, for all their support in access to established diagnostics.

## Contributions

FM, MB, and LU designed the study. LU conceptualised the study. FM conducted all laboratory work. TR conducted all computational analyses under LU’s supervision. KL assisted with laboratory optimisations. LW assisted with sequencing optimisations. SC, LK, and RS assisted with the established diagnostics. FM and LU wrote the manuscript, with input from all co-authors.

## Data and code availability

All genomic data will be made publicly available as bam files via ENA. All computational code is publicly available at https://github.com/ttmgr/GenomicsForOneHealth/tree/main/Food_Safety.

## Methods

### Bacterial strains and culture conditions

Study strains were selected to represent the genetic, ecological, and epidemiological diversity of *L. monocytogenes*, including lineage I (outbreak-associated), lineage II (environmentally persistent), and lineage III (low-virulence) strains (**Table 1**). Additional *Listeria* species (*L. innocua, L. ivanovii*, and *L. welshimeri*) were included as niche-overlapping controls to reflect naturally co-occurring *Listeria populations*.

**Table 1.** *Listeria* species strains.

| Strain ID | Description <sup>a</sup> | Source | Reference |
| --- | --- | --- | --- |
| ***L. monocytogenes*** |  |  |  |
| EGDe | LII, serotype 1/2a, ST35, CC9 | Reference strain | (Glaser et al., 2001) |
| N13-0119 | LII, serotype 1/2a, ST121, CC121 | Human listeriosis | (Althaus et al., 2014) |
| LL195 | LI, serotype 4b, ST1, CC1 | Swiss listeriosis outbreak | (Bille, 1990) |
| LMNC326 | LIII, serotype 4a/4c, ST70, CC70 | Ruminant listeriosis | (Dreyer et al., 2016) |
| ***Listeria* spp** |  |  |  |
| *L. innocua* J5051 | Control with niche overlap | Reference strain | (Guldimann et al., 2015) |
| *L. ivanovii* Nr26 | Control with niche overlap | Food processing plant | - |
| *L. welshimeri* Nr14 | Control with niche overlap | Vegetation | - |

\* L: Lineage, ST: Sequence type, CC: Clonal complex,

Background microbiota strains (**Table 2**) were selected to represent bacterial taxa commonly co-occurring with *L. monocytogenes* across food-processing environments, including meat, dairy, and fresh produce facilities, based on previous studies (Xu et al., 2023; Wagner et al., 2021; Barcenilla et al., 2024). Growth capacity in *Listeria* selective enrichment media was experimentally validated for all strains to ensure compatibility with enrichment conditions: Briefly, growth assays were performed in 96-well microplates by inoculating 20 µL of bacterial suspension (∼10^3^ CFU/mL) into 180 µL of HFB per well. Plates were incubated at 30°C with continuous shaking for 24 h. Optical density was measured every 30 min using a Synergy HT microplate reader (BioTek Instruments). Experiments were performed with three independent biological replicates, each measured in technical triplicate. The resulting growth patterns (**Table 2**) were consistent with previous findings (Wagner et al., 2021). This enabled a controlled and realistic assessment of microbial competition, enrichment-associated bias, and detection performance in complex communities.

**Table 2.** Background microbiota strains.

| Strain ID | Species | Potential growth in *Listeria* enrichment media |
| --- | --- | --- |
| W9 | *Enterococcus faecalis* | + |
| Nr.88 | *E. faecalis* | - |
| Nr.2 | *Bacillus cereus* | + |
| Wr3 | *B. subtilis* | - |
| W25 | *Pseudomonas aeruginosa* | - |
| Nr.36 | *P. aeruginosa* | + |
| W34 | *Staphylococcus aureus* | + |
| Nr.10 | *S. aureus* | + |
| W3 | *Escherichia coli* | - |
| BG1A | *Yersinia enterocolitica* | + |
| BG403 | *Y. enterocolitica* | - |
| N16-166 | *Y. frederiksenii* | + |
| Nr.9 | *Rhodococcus equi* | - |
| D 510-96 | *Proteus mirabilis* | - |

Bacteria were stored at −80°C in brain heart infusion medium (BHI, Oxoid) supplemented with 20% glycerol. Strains were initially grown overnight on BHI agar plates at 37°C to obtain single colonies, and then pre-cultured twice in 10-ml BHI broth (37°C, 150 rpm) for 18 h, to obtain stationary-phase secondary cultures. Unless otherwise stated, such secondary BHI pre-cultures were used for experiments. The overall experimental design, including sample spiking, enrichment strategy, and comparison of culture-based, direct metagenomic, and quasi-metagenomic workflows, is summarised in **Figure 1**.

### Preparation of inocula

Secondary stationary-phase BHI cultures prepared as described above were diluted in sterile physiological saline (0.9% NaCl) and standardised to approximately 1 × 10^9^ CFU/ml. Equal volumes of standardised cultures were combined to generate (i) a seven-strain *Listeria* spp. cocktail and (ii) a thirteen-strain background microbiota cocktail. The *Listeria* spp. cocktail (1 × 10^9^ CFU/ml) was serially diluted in tenfold steps to obtain working inocula ranging from 3 × 10^2^ to 3 × 10^8^ CFU/ml. The background microbiota cocktail was adjusted to 3 × 10^7^ or 3 × 10^8^ CFU/ml, depending on the experimental condition. In addition, to simulate non-growing environmental populations, *E. faecalis* Nr.88 (unable to grow in Half Fraser Broth (HFB)) was included as a non-growing background fraction at concentrations of 1 × 10^8^ or 1 × 10^9^ CFU/ml, for sponges and swabs, respectively.

Six final inocula were prepared, three for sponge samples and three for swab samples. For sponge swab inocula, *Listeria* spp. concentrations were adjusted to low (1 × 10^2^ CFU/mL; Lm2), medium (1 × 10^4^ CFU/ml; Lm4), and high (1 × 10^6^ CFU/ml; Lm6) levels, while maintaining a constant background microbiota concentration of 1 × 10^7^ CFU/ml (**Table 3**). For Zymo and cotton swab inocula, the same target concentrations (Lm2, Lm4, and Lm6) were used, but initial inoculum concentrations were increased by one log to compensate for the lower inoculation volume (100 µl versus 1 ml for sponge samples). Detailed inoculum compositions and mixing ratios are provided in **Supplementary Tables S5 and S6**.

**Table 3.** Target concentrations of *Listeria* spp. and background microbiota on swabs.

| Inoculum | Target *Listeria* spp (CFU/ml) | Background microbiota (CFU/ml) | Non-culturable background strain (CFU/ml) |
| --- | --- | --- | --- |
| Lm2 | $1 \times 10^2$ | $1 \times 10^7$ | $1 \times 10^8$ |
| Lm4 | $1 \times 10^4$ | $1 \times 10^7$ | $1 \times 10^8$ |
| Lm6 | $1 \times 10^6$ | $1 \times 10^7$ | $1 \times 10^8$ |

Actual inoculum concentrations were retrospectively confirmed by tenfold serial dilutions of samples in sterile saline and plating in duplicate on BHI agar (total viable counts) and ALOA (Agar *Listeria* according to Ottaviani and Agosti) agar for selective enumeration of *Listeria* spp. Plates were incubated at 30 °C (BHI) or 37 °C (selective media), and colonies were counted to determine CFU/ml.

### Sampling matrix spiking and processing

For direct DNA extraction, spiked sponge samples (Sponge-Stick with Letheen Broth, Neogen) were homogenised using a Stomacher 400 laboratory blender (Seward) for 7 min at medium speed. Following homogenization, sponges were incubated at 4 °C to allow dissipation of air bubbles and then manually compressed to recover the liquid fraction, yielding ∼4–6 ml of homogenate per sample. The homogenate was transferred to sterile tubes and centrifuged at 10,000 rpm for 20 min at 4 °C to pellet microbial biomass. Pellets were stored at −20 °C until further processing. DNA was extracted using the DNeasy PowerSoil Pro Kit (Qiagen) according to the manufacturer’s instructions. Spiked cotton and Zymo swabs were supplemented with 23 µl of DNA/RNA Shield (Zymo Research), cut to ∼3 cm in length, and transferred directly into ZR BashingBead™ Lysis Tubes (0.1 and 0.5 mm; Zymo Research). Samples were stored at −20 °C until further processing. DNA extraction was performed using the ZymoBIOMICS DNA Microprep Kit (Zymo Research) according to the manufacturer’s protocol. This workflow enabled recovery of biomass for direct metagenomic analysis while allowing comparison of matrix-dependent DNA retention effects.

Enrichment was performed for two distinct downstream applications: quasi-metagenomics, in which DNA was extracted from the mixed enrichment culture for community-level sequencing, and culture-based whole-genome sequencing, in which a single presumptive *L. monocytogenes* colony was isolated after enrichment and plated culture. Enrichment was only applied to sponges as the reference matrix for culture-based environmental surveillance. Quasi-metagenomic sampling time points (4 h, 12 h, and 24 h in HFB, and an additional 24 h in Fraser Broth (FB)) were selected to capture early, intermediate, and late stages of enrichment and to evaluate their impact on detection sensitivity and community composition. Sponges were supplemented with 90 mL HFB and homogenised using a stomacher for 7 min. The products were incubated statically at 30 °C for 24 h, and aliquots were collected at 4 h (9 ml), 12 h (4.5 ml), and 24 h (1.8 ml). Following 24 h of primary enrichment in HFB, 100 µl of each culture was transferred into 10 ml Fraser Broth (FB) and incubated at 37 °C for 48 h. Samples (1.8 ml) were collected at 24 h and 48 h. All samples were centrifuged at 13,000 rpm for 5 min at 4 °C, supernatants were discarded, and cell pellets were stored at −20 °C until further analysis. For cultures grown in FB, at 24 h and 48 h, a loopful was plated onto ALOA and PALCAM agar plates and incubated at 37 °C for 24–48 h. For the culture-based workflow, a single presumptive *L. monocytogenes* isolate was selected based on characteristic colony morphology and processed for whole-genome nanopore sequencing.

Each spiking experiment was run in two biological replicates.

### DNA extraction optimisations

Spiking experiments were performed using defined *L. monocytogenes* inocula in combination with complex background microbiota to simulate realistic environmental contamination scenarios. DNA extraction protocols were evaluated across sponges and two swab types. Extractions were performed using the DNeasy PowerSoil Pro Kit, DNeasy Blood & Tissue Kit (Qiagen), as well as the ZymoBIOMICS DNA Miniprep Kit and ZymoBIOMICS DNA Microprep Kit (Zymo Research). Performance was assessed by DNA yield and purity, retention effect, inhibitor carryover, and preservation of the microbial composition (data not shown).

In the final protocol, genomic DNA was extracted from cell pellets collected using either the DNeasy PowerSoil Pro Kit (sponge metagenomics), the ZymoBIOMICS DNA Microprep Kit (swab metagenomics, t_4_, t_12_), or the ZymoBIOMICS DNA Miniprep Kit (t_24_, t_48_), according to the manufacturers’ instructions. The ZymoBIOMICS DNA Microprep protocol was modified by omitting the Zymo-Spin III-F filter step and including an additional wash of the Zymo-Spin IC column with 200 µL of ZymoBIOMICS Wash Buffer 2. For all extractions, continuous bead beating was conducted at room temperature using a Vortex Genie (Scientific Industries) according to the manufacturer’s instructions. DNA concentration and purity were assessed using a NanoDrop spectrophotometer and the QuantiFluor ONE dsDNA System (Promega).

### Genomic approaches

Nanopore sequencing libraries were prepared using the Rapid Barcoding Kit 96 (SQK-RBK114.96) according to the manufacturer’s instructions per sample types: direct metagenomic samples, quasi-metagenomic samples collected at each enrichment time point, and isolate whole-genome sequencing samples derived from the culture-based workflow. Briefly, genomic DNA of each sample was subjected to transposase-mediated fragmentation and rapid barcode attachment, followed by pooling of barcoded samples. Two barcodes per sample were used for all low-concentration samples. Each library was sequenced on one MinION R10.4.1 flow cell (FLO-MIN114); For all metagenomic and quasi-metagenomic samples, adaptive sampling was applied to half of the flow cell using the *L. monocytogenes* EGDe reference genome NC_003210 as enrichment target.

### Computational analysis

The nanopore data were basecalled using Dorado SUP v5.0.0. Sequencing adapters and barcodes were removed using Chopper v0.11.0; sequencing reads shorter than 100 bp were filtered out with NanoFilt v2.8.0; and per-sample read-quality metrics (read count, N50, total bases, mean read quality) were computed using NanoStat v1.6.0.

Quality-filtered reads were mapped with minimap2 v2.30 using -x map-ont --secondary=no against a panel of sixteen bacterial genomes: four *L. monocytogenes* reference strains (EGDe, LL195, LMNC326, N13-0119); three non-*monocytogenes Listeria* species (*L. innocua* J5051, *L. ivanovii* Nr26, *L. welshimeri* Nr14); and nine environmental background species (*Bacillus cereus, B. subtilis, Escherichia coli, Pseudomonas aeruginosa, Proteus mirabilis, Rhodococcus equi, Staphylococcus aureus, Yersinia enterocolitica, Yersinia frederiksenii*). Per-strain statistics (mapped reads, coverage breadth, mean depth, mean MAPQ) were computed from samtools coverage output. *L. monocytogenes* recovery was quantified as the percentage of *L. monocytogenes* reads (and bases) out of the total per-sample read (and base) count from NanoStat.

Comparisons of adaptive sampling (AS) and normal (N) sequencing of quasi-metagenomic and metagenomic samples were performed using two-sided Mann–Whitney U tests. Raw p-values are reported without multiple testing adjustment. All analyses were performed in Python v3.12 using pandas, NumPy, and SciPy; all figures were generated using matplotlib and seaborn.

Biological replicates were treated as independent observations.

## References

Aalto-Araneda M, Pöntinen A, Pesonen M, Corander J, Markkula A, Tasara T, Stephan R, Korkeala H. Strain Variability of *Listeria monocytogenes* under NaCl Stress Elucidated by a High-Throughput Microbial Growth Data Assembly and Analysis Protocol. Appl Environ Microbiol. 2020 Mar 2;86(6):e02378–19. doi: 10.1128/AEM.02378-19. PMID: 31900307; PMCID: PMC7054083.

Allen JM. Rediscovering the wild: MiFoDB brings fermented food microbiomes into focus. mSystems. 2025 Sep 23;10(9):e0059925. doi: 10.1128/msystems.00599-25. Epub 2025 Aug 19. PMID: 40827869; PMCID: PMC12455962.

Barcenilla, C., Cobo-Díaz, J.F., De Filippis, F. et al. Improved sampling and DNA extraction procedures for microbiome analysis in food-processing environments. Nat Protoc 19, 1291–1310 (2024). 10.1038/s41596-023-00949-x

Barcenilla, C., Cobo-Díaz, J.F., Puente, A. et al. In-depth characterization of food and environmental microbiomes across different meat processing plants. Microbiome 12, 199 (2024). 10.1186/s40168-024-01856-3

Bille J. 1990. Epidemiology of human listeriosis in Europe with special reference to the Swiss outbreak, p 71–74 In Miller AJ, Smith JL, Somkuti GA, editors. Foodborne listeriosis. Elsevier, New York.

Bellankimath, A.B., Branders, S., Kegel, I. et al. Metagenomic sequencing enables accurate pathogen and antimicrobial susceptibility profiling in complicated UTIs in approximately four hours. Nat Commun 17, 187 (2026). 10.1038/s41467-025-66865-8

Bruhn JB, Vogel BF, Gram L. Bias in the *Listeria monocytogenes* enrichment procedure: lineage 2 strains outcompete lineage 1 strains in University of Vermont selective enrichments. Appl Environ Microbiol. 2005 Feb;71(2):961–7. doi: 10.1128/AEM.71.2.961-967.2005. PMID: 15691954; PMCID: PMC546678.

Buytaers FE, Saltykova A, Mattheus W, Verhaegen B, Roosens NHC, Vanneste K, Laisnez V, Hammami N, Pochet B, Cantaert V, Marchal K, Denayer S, De Keersmaecker SCJ. Application of a strain-level shotgun metagenomics approach on food samples: resolution of the source of a *Salmonella* food-borne outbreak. Microb Genom. 2021 Apr;7(4):000547. doi: 10.1099/mgen.0.000547. PMID: 33826490; PMCID: PMC8208685.

Buchanan, Robert L.; Gorris, Leon G.M.; Hayman, Melinda M.; Jackson, Timothy C.; Whiting, Richard C. (2017): A review of Listeria monocytogenes: An update on outbreaks, virulence, dose-response, ecology, and risk assessments. In 0956-7135 75, pp. 1–13. DOI: 10.1016/j.foodcont.2016.12.016.

Buytaers FE, Verhaegen B, Van Nieuwenhuysen T, Roosens NHC, Vanneste K, Marchal K, De Keersmaecker SCJ. Strain-level characterization of foodborne pathogens without culture enrichment for outbreak investigation using shotgun metagenomics facilitated with nanopore adaptive sampling. Front Microbiol. 2024 Mar 1;15:1330814. doi: 10.3389/fmicb.2024.1330814. PMID: 38495515; PMCID: PMC10940517.

Caffrey EB, Olm MR, Kothe CI, Wastyk HC, Evans JD, Sonnenburg JL. MiFoDB, a workflow for microbial food metagenomic characterization, enables high-resolution analysis of fermented food microbial dynamics. mSystems. 2025 Sep 23;10(9):e0014125. doi: 10.1128/msystems.00141-25. Epub 2025 Aug 19. PMID: 40827925; PMCID: PMC12456020.

Cornu M, Kalmokoff M, Flandrois JP. Modelling the competitive growth of *Listeria monocytogenes* and *Listeria innocua* in enrichment broths. Int J Food Microbiol. 2002 Mar;73(2-3):261–74. doi: 10.1016/s0168-1605(01)00658-4. PMID: 11934034

Dailey RC, Welch LJ, Hitchins AD, Smiley RD. Effect of *Listeria seeligeri* or *Listeria welshimeri* on *Listeria monocytogenes* detection in and recovery from buffered *Listeria* enrichment broth. Food Microbiol. 2015 Apr;46:528–534. doi: 10.1016/j.fm.2014.09.008.

Delikanli-Kiyak B, Yilmaz I, Guldas M. Can Metagenomic Analyses Be Used Effectively in Safe Food Production? Food Sci Nutr. 2025 Aug 7;13(8):e70772. doi: 10.1002/fsn3.70772. PMID: 40786823; PMCID: PMC12331879.

Dreyer M, Aguilar-Bultet L, Rupp S, Guldimann C, Stephan R, Schock A, Otter A, Schüpbach G, Brisse S, Lecuit M, Frey J, Oevermann A. 2016. *Listeria monocytogenes* sequence type 1 is predominant in ruminant rhombencephalitis. Sci Rep 6:36419. doi:10.1038/srep36419.

EFSA and ECDC (European Food Safety Authority and European Centre for Disease Prevention and Control), (2025). The European Union One Health 2024 Zoonoses Report. EFSA Journal, 23(12), e9759. 10.2903/j.efsa.2025.9759

Ercolini D. Exciting strain-level resolution studies of the food microbiome. Microb Biotechnol. 2017 Jan;10(1):54–56. doi: 10.1111/1751-7915.12593. Epub 2017 Jan 3. PMID: 28044418; PMCID: PMC5270722.

Finn, Lawrence; Onyeaka, Helen; O’Neill, Sally (2023): *Listeria monocytogenes* Biofilms in Food-Associated Environments: A Persistent Enigma. In Foods (Basel, Switzerland) 12 (18). DOI: 10.3390/foods12183339.

Flörl L, Meyer A, Bokulich NA. Exploring sub-species variation in food microbiomes: a roadmap to reveal hidden diversity and functional potential. Appl Environ Microbiol. 2025 May 21;91(5):e0052425. doi: 10.1128/aem.00524-25. Epub 2025 Apr 30. PMID: 40304520; PMCID: PMC12093984.

Ghiotto G, Zampieri G, Campanaro S, Treu L. Strain-resolved metagenomics approaches applied to biogas upgrading. Environ Res. 2024 Jan 1;240(Pt 2):117414. doi: 10.1016/j.envres.2023.117414. Epub 2023 Oct 16. PMID: 37852461.

Glaser P, Frangeul L, Buchrieser C, Rusniok C, Amend A, Baquero F, Berche P, Bloecker H, Brandt P, Chakraborty T, Charbit A, Chetouani F, Couvé E, Daruvar A de, Dehoux P, Domann E, Domínguez-Bernal G, Duchaud E, Durant L, Dussurget O, Entian KD, Fsihi H, García-del Portillo F, Garrido P, Gautier L, Goebel W, Gómez-López N, Hain T, Hauf J, Jackson D, Jones LM, Kaerst U, Kreft J, Kuhn M, Kunst F, Kurapkat G, Madueno E, Maitournam A, Vicente JM, Ng E, Nedjari H, Nordsiek G, Novella S, Pablos B de, Pérez-Diaz JC, Purcell R, Remmel B, Rose M, Schlueter T, Simoes N, Tierrez A, Vázquez-Boland JA, Voss H, Wehland J, Cossart P. 2001. Comparative genomics of *Listeria* species. Science 294:849–852. doi:10.1126/science.1063447.

Guldimann C, Bärtschi M, Frey J, Zurbriggen A, Seuberlich T, Oevermann A. 2015. Increased spread and replication efficiency of *Listeria monocytogenes* in organotypic brain-slices is related to multilocus variable number of tandem repeat analysis (MLVA) complex. BMC Microbiol 15:134. doi:10.1186/s12866-015-0454-0.

Jacobsen, C. N. 1999. The influence of commonly used selective agents on the growth of *Listeria monocytogenes*. Int. J. Food Microbiol. 50:221–226. 10.1016/S0168-1605(99)00101-4

Kocurek B, Ramachandran P, Grim CJ, Morin P, Howard L, Ottesen A, Timme R, Leonard SR, Rand H, Strain E, Tadesse D, Pettengill JB, Lacher DW, Mammel M, Jarvis KG. Application of quasimetagenomics methods to define microbial diversity and subtype *Listeria monocytogenes* in dairy and seafood production facilities. Microbiol Spectr. 2023 Dec 12;11(6):e0148223. doi: 10.1128/spectrum.01482-23. Epub 2023 Oct 9. PMID: 37812012; PMCID: PMC10714831.

Liu Y, Xu Y, Xu X, Chen X, Chen H, Zhang J, Ma J, Zhang W, Zhang R and Chen J (2023) Metagenomic identification of pathogens and antimicrobial-resistant genes in bacterial positive blood cultures by nanopore sequencing. Front. Cell. Infect. Microbiol. 13:1283094. doi: 10.3389/fcimb.2023.1283094

Manqele A, Gcebe N, Pierneef RE, Moerane R, Adesiyun AA. Identification of Listeria species and Multilocus Variable-Number Tandem Repeat Analysis (MLVA) Typing of *Listeria innocua* and *Listeria monocytogenes* Isolates from Cattle Farms and Beef and Beef-Based Products from Retail Outlets in Mpumalanga and North West Provinces, South Africa. Pathogens. 2023 Jan 15;12(1):147. doi: 10.3390/pathogens12010147. PMID: 36678495; PMCID: PMC9862459.

Martin S, Heavens D, Lan Y, Horsfield S, Clark MD, Leggett RM. Nanopore adaptive sampling: a tool for enrichment of low abundance species in metagenomic samples. Genome Biol. 2022 Jan 24;23(1):11. doi: 10.1186/s13059-021-02582-x. PMID: 35067223; PMCID: PMC8785595.

Maury MM, Tsai YH, Charlier C, Touchon M, Chenal-Francisque V, Leclercq A, Criscuolo A, Gaultier C, Roussel S, Brisabois A, Disson O, Rocha EPC, Brisse S, Lecuit M. Uncovering *Listeria monocytogenes* hypervirulence by harnessing its biodiversity. Nat Genet. 2016 Mar;48(3):308–313. doi: 10.1038/ng.3501. Epub 2016 Feb 1. Erratum in: Nat Genet. 2017 Mar 30;49(4):651. doi: 10.1038/ng0417-651b. Erratum in: Nat Genet. 2017 May 26;49(6):970. doi: 10.1038/ng0617-970d. PMID: 26829754; PMCID: PMC4768348.

Maury MM, Bracq-Dieye H, Huang L, Vales G, Lavina M, Thouvenot P, Disson O, Leclercq A, Brisse S, Lecuit M. Hypervirulent *Listeria monocytogenes* clones’ adaption to mammalian gut accounts for their association with dairy products. Nat Commun. 2019 Jun 6;10(1):2488. doi: 10.1038/s41467-019-10380-0. Erratum in: Nat Commun. 2019 Aug 6;10(1):3619. doi: 10.1038/s41467-019-11625-8. PMID: 31171794; PMCID: PMC6554400.

Milillo SR, Friedly EC, Saldivar JC, Muthaiyan A, O’Bryan C, Crandall PG, Johnson MG, Ricke SC. A review of the ecology, genomics, and stress response of *Listeria innocua* and *Listeria monocytogenes*. Crit Rev Food Sci Nutr. 2012;52(8):712–25. doi: 10.1080/10408398.2010.507909. PMID: 22591342.

Muchaamba F, Eshwar AK, Stevens MJA, Stephan R, Tasara T. Different Shades of *Listeria monocytogenes*: Strain, Serotype, and Lineage-Based Variability in Virulence and Stress Tolerance Profiles. Front Microbiol. 2022 Jan 4;12:792162. doi: 10.3389/fmicb.2021.792162. PMID: 35058906; PMCID: PMC8764371.

Nüesch-Inderbinen M, Horlbog JA, Cernela N, Stevens MJA, Stephan R. Whole-genome sequence-based surveillance of human clinical *Listeria monocytogenes* isolates from Switzerland, 2019–2024, Open Forum Infectious Diseases, 2026;, ofag222, 10.1093/ofid/ofag222

Olsen NS, Riber L. Metagenomics as a Transformative Tool for Antibiotic Resistance Surveillance: Highlighting the Impact of Mobile Genetic Elements with a Focus on the Complex Role of Phages. Antibiotics (Basel). 2025 Mar 12;14(3):296. doi: 10.3390/antibiotics14030296. PMID: 40149106; PMCID: PMC11939754.

Osek J, Lachtara B, Wieczorek K. *Listeria monocytogenes* - How This Pathogen Survives in Food-Production Environments? Front Microbiol. 2022 Apr 26;13:866462. doi: 10.3389/fmicb.2022.866462. PMID: 35558128; PMCID: PMC9087598.

Ottesen A, Ramachandran P, Chen Y, Brown E, Reed E, Strain E. Quasimetagenomic source tracking of *Listeria monocytogenes* from naturally contaminated ice cream. BMC Infect Dis. 2020 Jan 29;20(1):83. doi: 10.1186/s12879-019-4747-z. PMID: 31996135; PMCID: PMC6990534.

Painset A, Björkman JT, Kiil K, Guillier L, Mariet JF, Félix B, Amar C, Rotariu O, Roussel S, Perez-Reche F, Brisse S, Moura A, Lecuit M, Forbes K, Strachan N, Grant K, Møller-Nielsen E, Dallman TJ. LiSEQ - whole-genome sequencing of a cross-sectional survey of *Listeria monocytogenes* in ready-to-eat foods and human clinical cases in Europe. Microb Genom. 2019 Feb;5(2):e000257. doi: 10.1099/mgen.0.000257. Epub 2019 Feb 18. PMID: 30775964; PMCID: PMC6421348.

Parks DH, Newell RJP, Ginn AN, Bowerman KL, Alsheikh-Hussain A, Fang L, Shah S, MacDonald S, Wimpenny T, Evans P, Arias Guzman NE, Pribyl AL, Tyson GW, Hugenholtz P, Krause L, Newcombe J, Griffin P, Wehrhahn MC, Angel NZ, Wood DLA. Metagenomics enables parallel detection of 176 clinically relevant targets from faecal samples. Front Cell Infect Microbiol. 2026 Feb 23;16:1759322. doi: 10.3389/fcimb.2026.1759322. PMID: 41809988; PMCID: PMC12968262.

Quijada, N.M., Cobo-Díaz, J.F., Valentino, V. et al. The food-associated resistome is shaped by processing and production environments. Nat Microbiol 10, 1854–1867 (2025). 10.1038/s41564-025-02059-8

Scotter SL, Langton S, Lombard B, Schulten S, Nagelkerke N, In’t Veld PH, Rollier P, Lahellec C. Validation of ISO method 11290 part 1--detection of *Listeria monocytogenes* in foods. Int J Food Microbiol. 2001 Mar 20;64(3):295–306. doi: 10.1016/s0168-1605(00)00462-1. PMID: 11294351.

Segata N. On the Road to Strain-Resolved Comparative Metagenomics. mSystems. 2018 Mar 13;3(2):e00190–17. doi: 10.1128/mSystems.00190-17. PMID: 29556534; PMCID: PMC5850074.

Suh JH, Knabel SJ. Comparison of different enrichment broths and background flora for detection of heat-injured *Listeria monocytogenes* in whole milk. J Food Prot. 2001 Jan;64(1):30–6. doi: 10.4315/0362-028x-64.1.30. PMID: 11198438.

Tan CSC, et al., Characterising the microbial and antimicrobial resistance signatures of hospital-acquired pneumonia using nanopore metagenomic sequencing. bioRxiv 2025.08.09.669460; doi: 10.1101/2025.08.09.669460

Ürel H, Sauerborn E, Gebhardt F, Wantia N, Biggel M, Muchaamba F, Foster-Nyarko E, Brugger SD, Urban L. Nanopore metagenomic sequencing links clinically relevant resistance determinants to pathogens. doi: 10.64898/2026.02.16.706128

Wagner E, Fagerlund A, Langsrud S, Møretrø T, Jensen MR, Moen B. Surveillance of *Listeria monocytogenes*: Early Detection, Population Dynamics, and Quasimetagenomic Sequencing during Selective Enrichment. Appl Environ Microbiol. 2021 Nov 24;87(24):e0177421. doi: 10.1128/AEM.01774-21. Epub 2021 Oct 6. PMID: 34613762; PMCID: PMC8612253.

Walsh AM, Crispie F, Daari K, O’Sullivan O, Martin JC, Arthur CT, Claesson MJ, Scott KP, Cotter PD. Strain-Level Metagenomic Analysis of the Fermented Dairy Beverage Nunu Highlights Potential Food Safety Risks. Appl Environ Microbiol. 2017 Aug 1;83(16):e01144–17. doi: 10.1128/AEM.01144-17. PMID: 28625983; PMCID: PMC5541208.

Wang J, den Bakker HC, Denes TG. A critical review of metagenomic approaches for foodborne pathogen surveillance. Crit Rev Food Sci Nutr. 2025;65(33):8488–8501. doi: 10.1080/10408398.2025.2503453. Epub 2025 Jun 6. PMID: 40478725.

Xu ZS, Ju T, Yang X, Gänzle M. A Meta-Analysis of Bacterial Communities in Food Processing Facilities: Driving Forces for Assembly of Core and Accessory Microbiomes across Different Food Commodities. Microorganisms. 2023 Jun 14;11(6):1575. doi: 10.3390/microorganisms11061575. PMID: 37375077; PMCID: PMC10304558.

Zilelidou E, Karmiri CV, Zoumpopoulou G, Mavrogonatou E, Kletsas D, Tsakalidou E, Papadimitriou K, Drosinos E, Skandamis P. *Listeria monocytogenes* Strains Underrepresented during Selective Enrichment with an ISO Method Might Dominate during Passage through Simulated Gastric Fluid and In Vitro Infection of Caco-2 Cells. Appl Environ Microbiol. 2016 Dec 1;82(23):6846–6858. doi: 10.1128/AEM.02120-16. Epub 2016 Sep 16. PMID: 27637880; PMCID: PMC5103084.

Zitz U, Zunabovic M, Domig KJ, Wilrich PT, Kneifel W. Reduced detectability of *Listeria monocytogenes* in the presence of *Listeria innocua*. J Food Prot. 2011 Aug;74(8):1282–7. doi: 10.4315/0362-028X.JFP-11-045. PMID: 21819654.

